# Securing algal endosymbiont communities for reef-building corals

**DOI:** 10.1101/2022.06.14.495714

**Authors:** Jessica Bouwmeester, Jonathan Daly, Mariko Quinn, E. Michael Henley, Claire Lager, Riley Perry, Christopher A. Page, Mary Hagedorn

**Affiliations:** Center for Species Survival, Smithsonian Conservation Biology Institute, Front Royal, VA 22630; Hawaii Institute of Marine Biology, Kaneohe, Hawaii 96744

**Keywords:** cryopreservation, Symbiodiniaceae, *Cladocopium*, zooxanthellae, coral reefs, *Lobactis scutaria*, coral bleaching, climate change

## Abstract

Photosynthetic dinoflagellates that live in symbiosis with corals (family Symbiodiniaceae) are fundamental for the survival of coral reef ecosystems. During coral bleaching events, it is assumed that these symbionts remain available in the water column, in sediments, or are seeded from unbleached coral colonies. Yet, this hypothesis has not been verified and it remains unclear whether some diversity of Symbiodiniaceae may be lost in the process. Culture methods have been developed for some Symbiodiniaceae, but for the vast majority of these photosynthetic symbionts, known culture methods are not successful at maintaining them for extensive periods. For these unculturable symbionts, cryopreservation, which places cells and tissues in suspended animation for days to decades, offers the best hope for saving the biodiversity of these crucial coral partners. Some cryopreservation processes use slow freezing, but if the cells are sensitive to low temperatures, as is the case for Symbiodiniaceae, then rapid freezing, called vitrification, is needed. We here, tested two published vitrification protocols that had been designed for algal symbionts extracted from Hawaiian corals, but we were unable to recover living symbionts after vitrification and warming. Therefore, we report a successful optimisation of the former vitrification protocols, which we tested on algal symbionts freshly extracted from three Hawaiian coral species, the development of ultra-rapid laser-warming cryopreservation techniques for symbionts, and banking procedures for algal symbionts. We also present some successful uptake of cryopreserved algal symbionts by coral larvae, although at a low rate. It is unclear why the former vitrification protocols failed but we propose that it may have been related to thermal stress and bleaching events that occurred on several occasions throughout the Hawaiian Islands. Maintenance of biodiversity is essential for sustaining functional, productive ecosystems with the adaptability to effectively recover from disturbances. By successfully cryopreserving and banking coral symbionts, we provide a critically needed component for securing Symbiodiniaceae biodiversity into the future.

## Introduction

Coral reefs are threatened globally and it is estimated that almost one-third of reef-forming coral species are at risk of extinction due to climate change (IPBES 2019). Therefore, the need for innovative strategies to conserve and secure reef biodiversity is urgent (National Academies of Sciences Engineering and Medicine 2019). Critical to the survival of coral reefs is the mutualistic relationship between corals and their photosynthetic endosymbionts, dinoflagellate algae from the family Symbiodiniaceae. The algal symbionts provide their hosts with photosynthetic products, thus fulfilling most of the corals’ metabolic needs (Muscatine 1990). Most corals acquire their algal symbionts from the surrounding environment during early developmental stages, through horizontal transmission (Abrego et al. 2009; Baird et al. 2009; Nitschke et al. 2016). The exact thermal tolerance of the algal symbionts is species-specific (Howells et al. 2016) but overall, thermal stress can damage the photosynthetic machinery in the chloroplasts and cause the production of reactive oxygen species (Weis 2008). When this occurs, the symbiotic relationship between corals and their photosynthetic symbionts breaks down, resulting in coral bleaching.

The effects of climate change on corals are well documented (Hughes et al. 2017); however, the effect of mass coral bleaching on the biodiversity of Symbiodiniaceae is largely unknown. When outside their coral hosts, Symbiodiniaceae can be found in sediments, in the water column, and on the surface of macroalgae (Littman et al. 2008; Quigley et al. 2017; Fujise et al. 2021), but it is not clear whether these symbiont species in their free-living phase can tolerate prolonged exposure to increased water temperatures (but see Bellantuono et al. 2019). Additionally, it has been suggested that some Symbiodiniaceae species may be exclusively symbiotic (González-Pech et al. 2019), in which case these symbionts may not persist in the environment for extensive periods without the host species present (Nitschke et al. 2016; Thornhill et al. 2017). Extreme bleaching events that result in high coral mortality may therefore have a significant impact on Symbiodiniaceae communities, comparable to intense fires that sterilise forest soils and prevent rapid regrowth (Bowd et al. 2019). Therefore, it is essential to secure the biodiversity of coral-associated Symbiodiniaceae while diverse populations still exist. This can be accomplished through maintaining these algal symbionts in cultures (Fitt et al. 1981; Polne-Fuller 1991; Santos et al. 2001), or through cryopreservation (Santiago-Vázquez et al. 2007; Hagedorn and Carter 2015; Lin et al. 2019). To date, many Symbiodiniaceae strains that associate with corals have not yet been successfully cultured (Santos et al. 2001; Krueger and Gates 2012; LaJeunesse et al. 2018), and therefore, until effective culture protocols are developed, these strains are best secured through cryopreservation—the storage of material at ultra-low temperatures—which can maintain living cells, tissues, and germplasm in a frozen state indefinitely. Cryopreservation and biobanking of coral sperm have been ongoing since 2008 in a variety of coral ecosystems (Hagedorn et al. 2012a; Hagedorn et al. 2012b; Hagedorn et al. 2019) and a similar effort to preserve Symbiodiniaceae would be highly beneficial to reef-building corals, which heavily rely on these algal symbionts for their survival.

Cryopreservation techniques for algal symbionts have been developed for cultured strains but post-thaw culture expansion time from these cryopreserved symbionts generally takes months (Chong et al. 2016; Lin et al. 2019), suggesting that the cryopreservation process for these cells may not yet be optimal (but see Kihika et al. 2022). Better symbiont cryopreservation methods would be highly beneficial for these cultured species, but mostly, they are critical for a vast range of coral-associated algal symbionts that are currently not culturable outside of their hosts (Krueger and Gates 2012; Voolstra et al. 2021). To date, cryopreservation of Symbiodiniaceae still remains challenging due to high sensitivity to cryoprotectants and to chilling (Hagedorn et al. 2010), even though some successful cryopreservation protocols have been reported with different measures of post-thaw viability for culturable symbionts (Chong et al. 2016; Lin et al. 2019; Kihika et al. 2022) as well as freshly extracted unculturable symbionts (Hagedorn and Carter 2015).

For this work, we used extracted algal symbiont communities from the genus *Cladocopium* (Symbiodiniaceae), which dominate the Hawaiian coral species *Lobactis scutaria, Porites compressa, and Leptastrea purpurea* (LaJeunesse et al. 2004). Here, we test previous cryopreservation protocols developed for unculturable *Cladocopium* symbionts and present a novel cryopreservation protocol for algal symbionts using ultra-rapid laser-warming cryopreservation techniques. We detail banking procedures for cryopreserved Symbiodiniaceae and finally, we test the inoculation of coral larvae with symbionts cryopreserved using the novel cryopreservation protocol.

## Materials and Methods

### Collection and Maintenance of Coral

Whole individuals of *Lobactis scutaria*, and 15 cm fragments of *P. compressa* and *Leptastrea purpurea* were collected from reef flats (1-2m depth) and reef slopes (2-5m depth) in Kaneohe Bay, Hawaii from 2018 to 2021 under permits to the Hawaii Institute of Marine Biology SAP 2019-16, SAP 2020-25, and SAP 2021-33, from the State of Hawaii Department of Land & Natural Resources. Corals chosen for collection were at least 10m apart from each other and were sampled from multiple reefs to ensure as much genetic diversity as possible and avoid collecting corals that shared the same genotype. Corals were used immediately after collection or maintained in free-flowing seawater aquarium systems connected directly to Kaneohe Bay with natural temperature and light exposure throughout the study.

### Isolation of Symbiodiniaceae

Symbiodiniaceae communities were freshly isolated from live coral tissue following Bouwmeester et al. (2022a) to obtain a clean and homogenous suspension of algal symbionts. The symbionts were then kept overnight in an antibiotic cocktail (Polne-Fuller 1991) and the next morning they were rinsed, passed through a 20 μm cell strainer to remove clumps, and concentrated to ~ 4 × 10^7^ cells/ml (Bouwmeester et al. 2022b). A fresh algal symbiont isolation was performed for each cryopreservation attempt.

### Vitrification and Thawing

Three vitrification protocols were tested (see Table 1): *P1* – droplets of algal symbionts in solution following Hagedorn and Carter (2015), *P2* – droplets of algal symbionts encapsulated in agar following Hagedorn and Carter (2015), and *P3* – droplets of algal symbionts encapsulated in alginate (this paper; electronic supplemental material). Two thawing methods were tested: convective warming in 30 °C FSW following Hagedorn & Carter (2015), and laser warming using an infrared laser following Khosla *et al*. (2017) with modifications as described in Daly *et al*. (2018). Laser warming has shown to significantly improve the cryopreservation of complex cells and tissues (Jin et al. 2014), such as fish embryos (Khosla et al. 2017) and coral larvae (Daly et al. 2018), which are large, lipid-filled and have complex membrane permeabilities. The details of the *P3* vitrification protocol and the laser-warming methods are detailed in the electronic supplemental material.

**Table 1:** The three vitrification protocols (*P1*, *P2* and *P3*) tested for Symbiodiniaceae

| Protocol description | *P1*: Vitrification of algal symbionts in solution following (Hagedorn and Carter 2015) | *P2*: Vitrification of algal symbionts encapsulated in agar following (Hagedorn and Carter 2015) | *P3*: Vitrification of algal symbionts encapsulated in alginate (this paper) |
| --- | --- | --- | --- |
| Initial medium containing Symbiodiniaceae | *P1*: FSW | *P2*: 1.5% agar in FSW | *P3*: 2% alginate in deionised water with 1M trehalose |
| Step 1 | *P1, P2, P3*: 60 minutes in 10% DMSO in FSW |  |  |
| Step 2 | *P1, P2, P3*: 30 minutes in 0.5M trehalose and 10% DMSO in FSW |  |  |
| Step 3 | *P1, P2*: 5 minutes in 0.5M trehalose, 10% DMSO and 5% MeOH in FSW |  | *P3*: 3 minutes in 0.5M trehalose, 10% DMSO and 5% MeOH in FSW |
| Step 4 | *P1, P2*: N/A |  | *P3*: 30 seconds in DPT vitrification solution (1M trehalose, 10% DMSO, 5% PG in PBS) |

### Assessments of Symbiodiniaceae Viability

The main role of algal symbionts when in symbiosis with their coral host is to provide essential photosynthetic products to their host, covering the coral’s energetic needs for growth and reproduction. We used a Junior Pulse Amplitude Modulated fluorometer (Junior-PAM, Walz, Germany) to obtain the photosynthetic yield, an indication of the viability of photosynthesis in the Photosystem II (for more details, see electronic supplemental material). Photosystem II yield values were paired with light microscope observations (Olympus BX-41), to verify the absence of any other photosynthetic organisms and to assess the morphology of the post-thaw cells.

Four treatments were tested and assessed for each isolated Symbiodiniaceae community, (i) Cryo/Cryopreserved = symbionts encapsulated in alginate, exposed to vitrification solutions, vitrified and laser warmed; (ii) Tox/Toxicity = a toxicity control with symbionts encapsulated in alginate, exposed to vitrification solutions only; (iii) Live = a live control with symbionts encapsulated in alginate only; and, (iv) Dead = dead control, with symbionts encapsulated in alginate, exposed to vitrification solutions, vitrified and slow-warmed or with symbionts exposed to multiple freeze-thaw cycles.

### Production of Coral Larvae

Coral (*L. scutaria*) individuals (n = 30-50) were placed in separate bowls at 16:00 one to three days after the full moon during the 2021 summer months, covering the known times of spawning of that species in Hawaii (Hagedorn et al. 2016). Between 16:30 and 18:30, the corals began to spawn by expelling brief puffs of eggs or sperm. Eggs were collected from each female bowl with a transfer pipette, and transferred to clean bowls. Sperm was captured with a transfer pipette directly from around the mouth of the fungiids as sperm was being released. Each night, the sperm from 5-7 individuals was pooled, and the sperm motility and concentration were assessed with Computer Assisted Sperm Analysis following Zuchowicz et al. (2021). The fresh sperm was added into the egg bowls for a final ratio of approximately 1:10000 (eggs:sperm) and was allowed to fertilise for 1 hour. The water from the egg bowls was then gently replaced to dilute and remove as much sperm as possible, and the eggs were left to develop in a 26°C environment. Daily cleaning with filtered seawater maintained the larvae in good health.

### Inoculation of Coral Larvae with Cryopreserved Symbionts

Larvae from the coral *L. scutaria* are most likely to take up their algal symbionts three to four days after fertilisation (Schwarz et al. 1999). We conducted independent symbiont uptake trials on both larval development days. Symbiont uptake was tested using the following treatments: (i) Freshly isolated algae unfiltered= freshly isolated algae that were roughly cleaned and that were not filtered through 70, 40, or 20 μm mesh filters, therefore retaining some coral cells and debris (n=9); (ii) Freshly isolated algae clean= thoroughly cleaned of most host coral cells and embedded in alginate (n=3); (iii) Cryopreserved algae= symbionts thoroughly cleaned of most host coral cells and embedded in alginate, vitrified and laser-warmed (n=45); (iv) No algae= no symbionts added (n=23); and, (v) Dead algae= symbionts cleaned of most host coral cells and embedded in alginate, vitrified and ambiently warmed (n=12) or symbionts exposed to several freeze-thaw cycles (n=3). One hundred to 200 larvae were used for each treatment, after exposure to filtered *Artemia* sp. to induce feeding behaviour (Schwarz et al. 1999). They were exposed to each treatment for 2 hours and then transferred to clean filtered seawater in new dishes. All inoculations with algal symbionts were conducted in 1-ml volumes. Inoculations with cryopreservation algal symbionts were tested at four different concentrations to determine the ideal concentration for successful uptake (i.e. 1.5 × 10^5^ cell/ml, 7.5 × 10^4^ cell/ml, 3 × 10^4^ cell/ml or 1.5 × 10^4^ cell/ml). All other inoculations were conducted at 1.5 × 10^5^ cell/ml. Larvae were then moved to 10-ml dishes and cleaned daily with filtered seawater until assessments 72 h later (except for the last batch of day 4 infections, which were assessed 48 h later). All algal symbiont uptake assessments were conducted under a compound microscope (Olympus BX-41) with a 10X objective. From each genotype and treatment, at least 100 larvae were transferred to a slide, immobilised with a coverslip and scored for symbiont uptake.

### Statistical Analysis

One-way ANOVAs were used to test for differences in symbiont photosynthetic yield rates among the cryopreserved, live control, toxicity control, and dead control treatments. Normality of the residuals was verified by plotting a histogram of the residuals against a normal distribution curve and homoscedasticity was verified by plotting the model residuals against the fitted model. Where relevant, post hoc tests were conducted with least square means pairwise comparisons with Tukey adjustment for multiple comparisons. Simple logistic regressions were used to test for differences in symbiont uptake success among the Cryopreserved, Live, No Symbiont, and Dead treatments. Where relevant, significance of each treatment were assessed using linear model contrasts. All statistical analyses were conducted using R (R Core Team 2019), and the R packages car (Fox and Weisberg 2019), rcompanion (Mangiafico 2019), lsmeans (Lenth 2016), ggplot2 (Wickham 2016), multcompView (Graves et al. 2015), and Rmisc (Hope 2013).

## Results

### Testing previously established cryopreservation protocols

We tested the two protocols that had been developed by Hagedorn and Carter (Hagedorn and Carter 2015), i.e. with and without encapsulation in agar. Despite repeated attempts, the algae’s photosynthetic activity post-cryopreservation, determined with a Junior-PAM, remained below 0.050, indicating that the cells were not functional. During these attempts, several issues were identified. First of all, the encapsulation in agar, which requires briefly mixing the algal symbionts with melted agar at ~60°C, was detrimental to the symbionts, causing a 50% drop in the photosynthetic health of the symbionts. Second, the algal symbionts were sensitive to the toxicity of the vitrification solutes, as measured by a further drop in the photosynthetic health of the symbionts. Third, the vitrification of symbionts was inconsistent, as some samples appeared clear while others were hazy, suggesting ice formation. Finally, less than a day post-cryopreservation, the post-thaw algal symbionts were swarming with bacteria, which likely further compromised recovery. The ultimate outcome was a year of post-thaw assessments that yielded no clearly viable symbionts, as assessed with a PAM fluorometer.

### Optimisation of algal symbiont cryopreservation

A series of modifications were made to optimise the cryopreservation protocol for Symbiodiniaceae freshly extracted from their coral hosts (for details and for the full protocol, see electronic supplemental material). Key changes that led to success were (i) enhanced cleaning of algal symbiont cells, removing most coral host cells; (ii) treating symbionts with antibiotics prior to cryopreservation; (iii) removal of salts from all solutions to reduce potential ice nucleation events; (iv) replacement of high temperature agar with low-temperature alginate; (v) improved equilibration and vitrification steps (Table 1); and (vi) ultra-rapid laser warming to reduce lethal intracellular ice.

### Post-thaw Viability Measurements

The optimised symbiont vitrification protocol was tested on three algal symbiont communities, each extracted from a different coral species from Kaneohe Bay, Hawaii, and their photosynthetic health was assessed 24 hrs later to determine their survival (Fig. 1, Table S1). Symbionts extracted from the coral *L. scutaria* had the highest functional photosynthetic rates post-thaw after the Cryopreservation treatment, retaining 73% photosynthetic function overall in comparison to the Live treatment (Fig. 1A), whereas cryopreserved *P. compressa* symbionts recovered 50% (Fig. 1B) and cryopreserved *L. purpurea* symbionts recovered 30% photosynthetic function overall in comparison to the Live treatment (Fig. 1C).

**Fig. 1.**
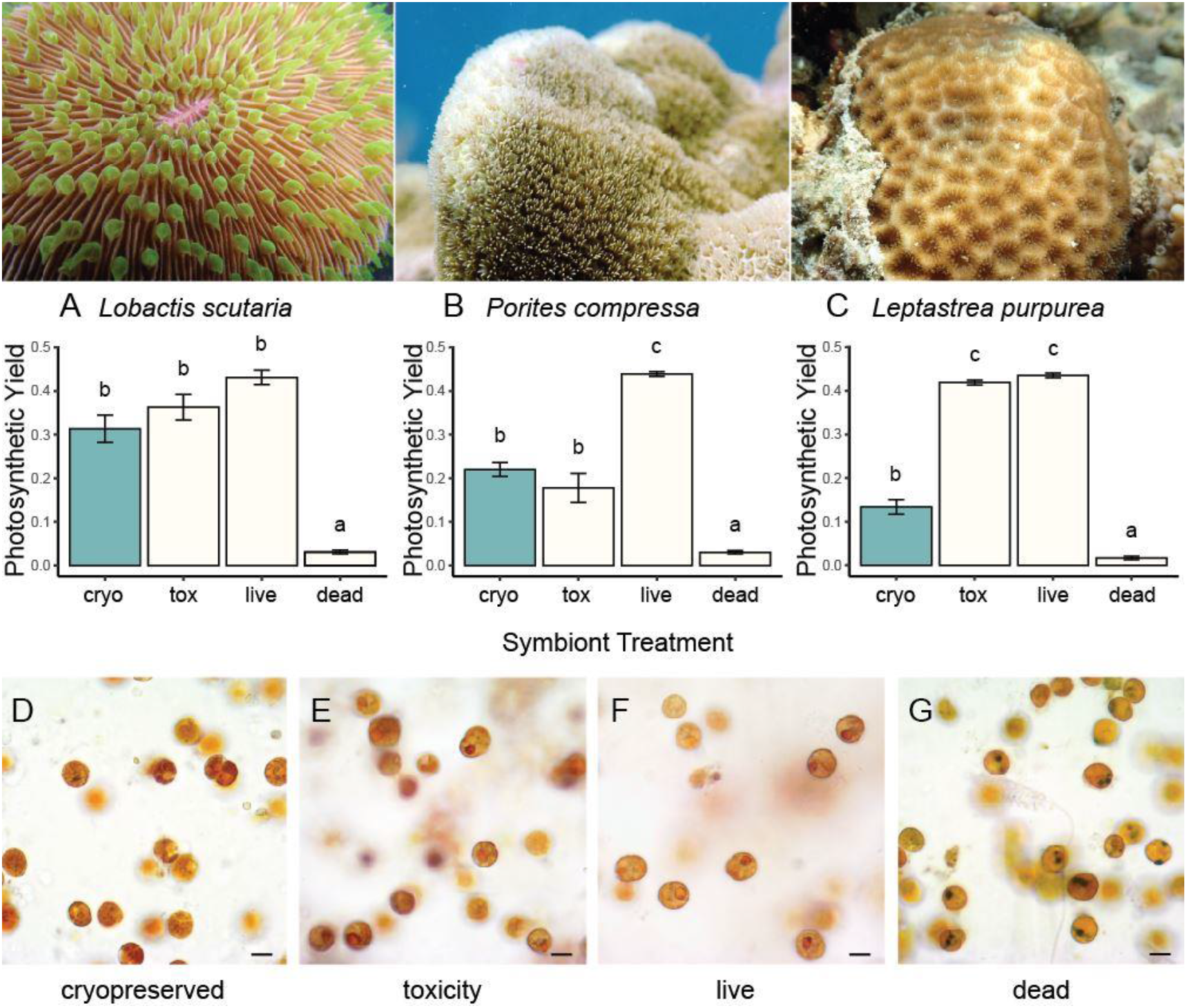
Photosynthetic yield and microscopy assessments of algal symbiont communities extracted from three Hawaiian coral species: (A) *Lobactis scutaria*, (B) *Porites compressa*, and (C) *Leptastrea purpurea*. The upper panel shows representative images of the host coral species. (A – C) Photosynthetic yield was representative of the health of the algal symbionts before and after cryopreservation. Four treatments were assessed for each species, (i) Cryo/Cryopreserved = symbionts encapsulated in alginate, exposed to vitrification solutions, vitrified and laser warmed (*L. scutaria*: n=13; *P. compressa*: n=17: *L. purpurea*: n=5); (ii) Tox/Toxicity = a toxicity control with symbionts encapsulated in alginate, exposed to vitrification solutions only (*L. scutaria*: n=5; *P. compressa*: n=3: *L. purpurea*: n=5); (iii) Live = a live control with symbionts encapsulated in alginate only (*L. scutaria*: n=5; *P. compressa*: n=4: *L. purpurea*: n=5); and, (iv) Dead = dead control, with symbionts encapsulated in alginate, exposed to vitrification solutions, vitrified and ambiently-warmed (*L. scutaria*: n=5; *P. compressa*: n=3: *L. purpurea*: n=6). In the Cryopreserved, Toxicity and Live treatments, viable symbionts with high photosynthetic yields were present. In contrast, the Dead symbiont treatment demonstrated little photosynthetic yield. Treatments that share the same lower-case letter in (A), (B), (C) are not significantly different from each other (one way ANOVA post hoc tests with least square means pairwise comparisons and Tukey adjustment for multiple comparisons; electronic supplemental material, Table S3). Data are presented as means ± SEM. Additionally, microscopic assessments of the algal symbionts 24 h post-treatment demonstrated no visible change in morphology in symbionts from the (D – F) Cryopreserved, Toxicity and Live treatments. However, the Dead symbiont treatments demonstrated degraded morphology. (G) Dead symbiont treatment. Note orange nuclei in most symbionts in (D – F), but few in (G). Scale bar = 10 μm.

These photosynthetic yields were paired with microscopic observations, to examine morphological changes in the cells and to rule out the presence of possible contaminating photosynthetic organisms. We observed that the symbionts from the Cryopreserved, Toxicity and Live treatments all shared a similar morphology. Specifically, these living cells exhibited a deep brown pigment, a spherical shape with smooth membranes and a distinguishable orange nucleus (Fig. 1 D – F). In contrast, symbionts from the Dead treatments shared another type of morphology. These dead symbionts had a lighter colour, a flatter, irregular shape with rough membranes and many had condensed, black nuclei (Fig. 1 G).

### Banking

Post-bleaching coral symbionts were not only vitrified successfully, but they were also banked successfully. Briefly, the vitrified symbionts encapsulated in alginate on the blades were immersed in liquid nitrogen, the blades removed from their holder, placed individually into pre-chilled cryovials and transferred to a cryobank where they were held at −190°C. This banking process started in April 2021 with symbionts extracted from the coral *L. scutaria* and was ongoing throughout the spring and summer. To use the samples, in our case for larval infection experiments, the blades were returned to their holders under liquid nitrogen and then were laser-warmed 24 h before their use. At 24 h post-thaw, the photosynthetic yield values of all the cryopreserved material banked ranged from 0.180 to 0.401 throughout this entire banking period, indicating 42-93% photosynthetic ability compared to control symbionts that had not undergone any treatment.

### Inoculation of L. scutaria coral larvae with cryopreserved algal symbionts

The uptake rate of dead symbionts killed by ambient warming was not different from those killed by repetitive freezing and warming (logistic regression, p = 0.313). Therefore, these groups were pooled into a single Dead algae treatment. The photosynthetic yields of the symbionts from the Freshly isolated algae unfiltered, Freshly isolated algal clean and Cryopreserved algae treatments demonstrated activity of Photosystem II, whereas the Dead treatments did not (electronic supplemental material, Table S2).

The Freshly isolated algae unfiltered treatment yielded twice as many larvae with symbionts than the Freshly isolated algae clean treatment (logistic regression, p < 2.2 × 10^−16^; Fig. 2A; electronic supplemental material, Table S3). The symbiont uptake success for the larvae exposed to the Cryopreserved algae treatment varied between 0 and 2.9 %, and was significantly higher than the No algae and Dead algae treatments (logistic regression, contrasts, p = 0.021; Fig. 2B; electronic supplemental material, Table S3). However, there was no difference between the Dead and No Symbionts treatments (logistic regression, contrasts, p = 0.720 and p = 0.726, respectively). Although four different concentrations of symbionts were used in Cryopreserved treatments (i.e., 1.5 × 10^5^, 7.5 × 10^4^, 3 × 10^4^ and 1.5 × 10^4^ cells/mL), the larval uptake rates of these different concentrations did not change (logistic regression, p = 0.846; electronic supplemental material, Table S3), and therefore these data were pooled into a single Cryopreserved algae treatment. The number of symbionts taken up by each larva was variable, with larvae containing 0 to 49 symbionts in the Live with Coral Cells treatment, 0 to 27 symbionts in the Live Clean treatment, 0 to 14 symbionts in the cryopreserved treatment, and 0 to 2 symbionts in the No Symbionts added treatment representing background symbiont uptake levels (electronic supplemental material, Fig. S1).

**Fig. 2.**
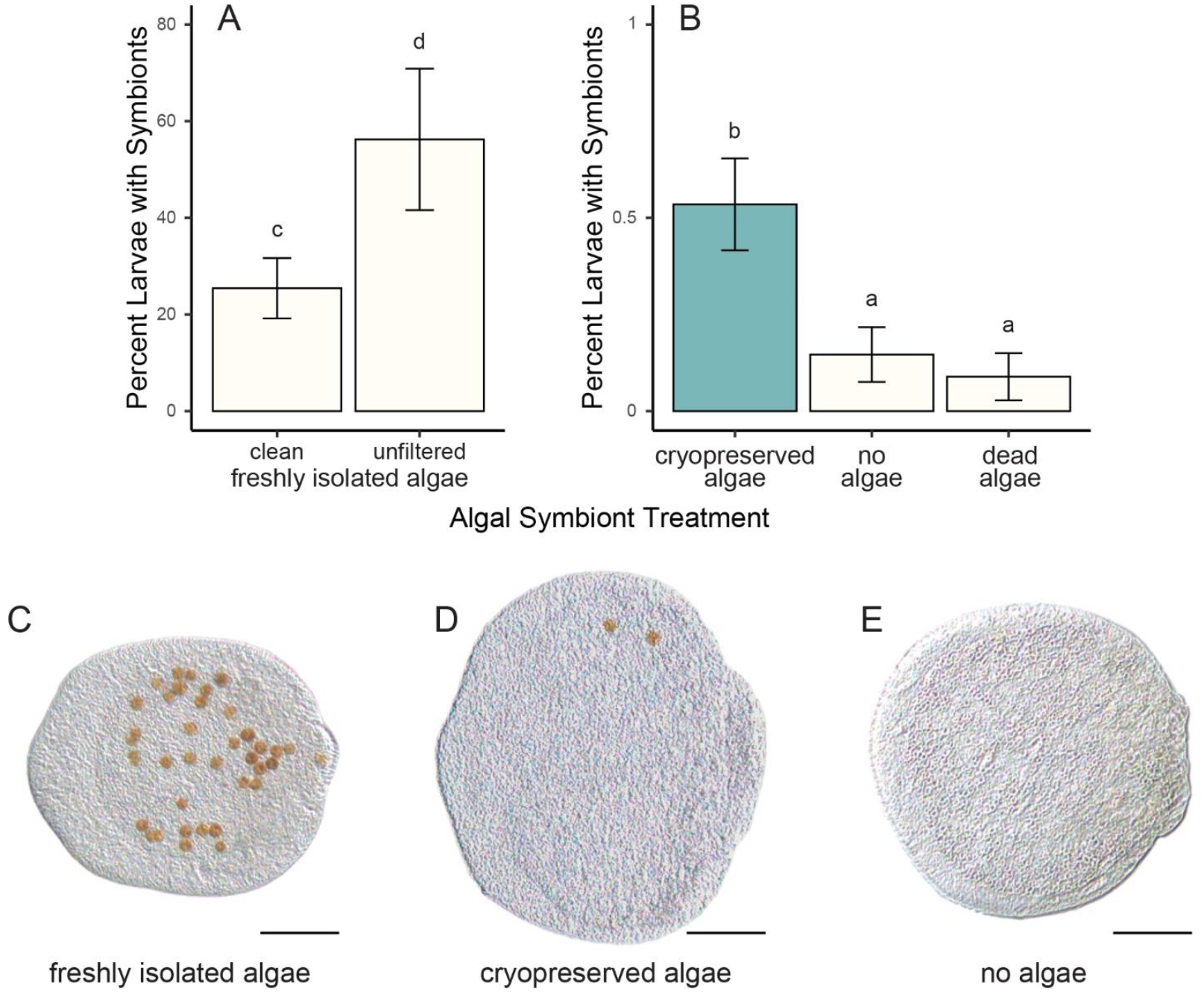
Symbiont uptake success for *Lobactis scutaria* larvae. (A) A comparison of the uptake success of the larvae exposed to symbionts from the two Freshly isolated algae treatments, clean (n=9) or unfiltered (n=3), which produced different infection success (logistic regression, p-value < 2.2 × 10^−16^). (B) The uptake success of the larvae exposed to the Cryopreserved algae (n=45, includes four algal symbiont concentrations that yielded similar uptake rates), No algae added (n=23), and Dead algae treatments (n=15). The larvae from the Cryopreserved treatment contained significantly more larvae than the larvae from the No algae and the Dead algae treatments (logistic regression, contrasts, p-value = 0.021). Treatments that share the same lower-case letter in (A), (B) are not significantly different from each other (logistic regression with posthoc linear model contrasts). Data are presented as means ± SEM. (C – D) Whole-mount larval light microscopic images of successful larval infections with (C) Live, here with coral cells, and (D) Cryopreserved symbiont treatments. There was no visible morphological difference between these two treatments. (E) Light microscopic image of a larva that did not absorb symbionts. All infection successes were measured 48 to 72 h post-treatment. Scale bar = 50 μm.

## Discussion

As we face ever-warming summers around the world, coral bleaching has the potential to reduce coral abundance and diversity. Therefore, forming a robust, comprehensive cryobank of the world’s algal symbionts is critical for ensuring the maintenance of global coral diversity throughout the next decades. We show that this is now possible with freshly isolated algal symbionts, with high survival rates post-cryopreservation.

The optimised Symbiodiniaceae cryopreservation protocol presented here successfully vitrified and laser-warmed algal symbionts communities isolated from three Hawaiian coral species. However, the failure of the protocol developed by Hagedorn and Carter (2015) was surprising, despite several attempts both with and without encapsulation of the symbionts in agar, given the Symbiodiniaceae were isolated from corals from the same region (i.e. Kaneohe Bay, Hawaii) and given all experiments were conducted in the same laboratory at the Hawaiian Institute of Marine Biology. This strongly suggests that the Symbiodiniaceae communities have changed between the last algal symbiont cryopreservation trials conducted by Hagedorn and Carter (2015) and the beginning of this work in 2018. Some Symbiodiniaceae physiology assessments between the same two periods indicated differences in chilling sensitivity, in cryoprotectant solution toxicity, and in membrane permeability (Bouwmeester et al. 2019). Such changes may be caused by a shift in the Symbiodiniaceae composition of communities isolated from corals in Kaneohe Bay. However, this shift would have had to occur in at least three coral species and remain stable over time. Alternatively, the physiology of the Symbiodiniaceae may have changed. Specifically, the composition of the cell and thylakoid membranes may have been altered, potentially, in response to climate change, progressively warming seawater temperatures around the main Hawaiian Islands and several coral bleaching events (Bahr et al. 2017; Couch et al. 2017; Rodgers et al. 2017; Ritson-Williams and Gates 2020). Similar membrane changes have indeed been experimentally observed in other studies that involved exposing algal symbionts to warmer temperatures (Tchernov et al. 2004; Díaz-Almeyda et al. 2011; Beltran et al. 2021).

In addition to changes associated with climate change, some species-specific adjustments to the cryopreservation protocol may need to be incorporated in the future to recover symbionts with different rates of post-thaw photosynthetic function. For example, post-cryopreservation photosynthetic function recovery was variable within symbionts from *L. scutaria* (78%) and *L. purpurea* (30%), potentially due to differences in toxicity, for which the cryopreservation protocol can be adapted. Interestingly, LaJeunesse et al. (2004) determined that both coral species in Hawaii shared the same dominant group of *Cladocopium* clade C1f symbionts in 2004, but the data, shown here, indicated that there was a physiological difference in their response to cryopreservation. This raises the question about whether they still host the same dominant symbionts, or whether other symbiont clades present (not detected in the above study), might have become more dominant. Therefore, further molecular work focused on genotyping current symbiont communities in the corals studied here would be highly informative. Alternatively, it is unknown whether cryopreservation could have selected some of the symbiont species/taxa in each symbiont community. It is common for coral species to host a diverse symbiont community (Mieog et al. 2007; Hume et al. 2015; Rouzé et al. 2017), and cryopreservation might have successfully cryopreserved one species/taxon over another, accidently selecting the one that is the most tolerant to cryopreservation. Similar selective processes have been identified in symbionts that are maintained in culture conditions, whereby the cultured symbiont species may not well reflect the original population from which it was extracted (Santos et al. 2001). Additional molecular work is therefore needed to confirm how well the cryopreserved symbiont communities match the original symbiont communities in each host coral species and whether selective changes occurred during the cryopreservation process.

Our data revealed low rates but successful inoculation of coral larvae with vitrified and laser-warmed algal symbionts. The successful uptake of cryopreserved symbionts and the stability of the symbiosis after three days in our experiments suggests that the cryopreserved symbionts retained sufficient integrity and function for recognition, uptake, and retention by the coral larvae. However, additional work is still needed to increase these rates. This study uses Symbiodiniaceae encapsulated in alginate but it is unclear whether the alginate may have detrimental effects on the initiation or the maintenance of symbiosis between the corals and their algae. Additionally, we found that freshly isolated symbionts, unfiltered and in solution (in FSW) yielded higher uptake rates than freshly isolated symbionts that were thoroughly cleaned and encapsulated in alginate. The reason for these higher uptake rates is not yet clear but identifying the factors that facilitate algal symbiont uptake may reveal to be highly information for cryopreserved algal symbionts.

Symbiodiniaceae cryopreservation is key to the functionality of coral bio-repositories as effective restoration tools, especially for endangered corals, such as the Caribbean coral *Dendrogyra cylindrus*, that may have limited selection of algal symbiont types (Lewis et al. 2019) and for corals that shuffle their symbionts due to seasonal changes and after massive bleaching (Hume et al. 2015; Lewis et al. 2019). Another important consideration is to understand the development of the complement of symbionts needed during early coral development. For example, some coral larvae have a complement of juvenile symbionts that shifts to an adult form later in life, depending on the environment (Koerner 2019). With the establishment of a successful method for the cryopreservation of coral symbionts of the family Symbiodiniaceae, future work will require scaling up the process for restoration-ready usage. This will require engaging with engineers to make this complex protocol and associated processes easier and faster, for overall increased efficiency and output (Tiersch 2011).

We cannot help save coral reefs through *ex situ* conservation means without equally balanced and robust frozen bio-repositories for both coral and algal symbionts. This study revealed that the inoculation of coral larvae with algal symbionts was possible with cryopreserved symbionts, opening new avenues for restoration of our reefs.

## Supporting information

electronic supplemental material

## Acknowledgements

The authors are grateful to Dr. Manuel Aranda (KAUST, Saudi Arabia) for providing feedback to this manuscript. Collection was performed with the appropriate permits from the state of Hawaii’s Department of Land and Natural Resources (Special Activity Permit # SAP 2018: 2011-1, 2019: 2012-63, 2020: 2013-47 and 2021: 2015-17). No institutional ethical approval was required for any of the experimental research described herein; however, animals were maintained with the highest husbandry standards. This work was supported by William H. Donner Family Foundation, the Paul M. Angell Family Foundation, the Volgenau Foundation, the Barrett Family Foundation, the Skippy Frank Foundation, the Compton Foundation, the Cedar Hill Foundation and the Anela Kolohe Foundation, the Smithsonian Conservation Biology Institute, the Smithsonian Women’s Committee and the Hawaii Institute of Marine Biology. Funders did not influence the interpretation of the project data or the decision to publish the findings.

## Notes

### Competing Interest Statement

The authors have declared no competing interest.

