## Supplementary material for "Securing algal endosymbiont communities for reef-building corals": electronic supplemental material

### Title: Securing Algal Symbiont Communities for Reef-Building Corals

**This PDF file includes:**

Supplementary Information: Optimisation of algal symbiont cryopreservation
Supplementary Information: Detailed laser-warming
Supplementary Information: Pulse amplitude modulated fluorometry
Supplementary Information: Optimised Protocol, Step-by-Step
Supplementary Information: References
Supplementary Figure S1
Supplementary Tables S1 to S3

### Supplementary Information: Optimisation of algal symbiont cryopreservation

Based on Pulse Amplitude Modulated fluorometry health assessments throughout each step of the cryopreservation protocol, we made a series of adjustments to the Symbiodiniaceae cryopreservation protocol. We encapsulated the symbionts in alginate as the symbionts needed to be encapsulated to minimise toxicity issues but did not survive well the encapsulation in heated agar. To improve vitrification of the sample, we started the protocol with spinning the symbionts in a centrifuge and resuspending them in deionised water with 1M trehalose sugar instead of filtered seawater. This step maintained iso-osmotic conditions but minimised the presence of salts that can become centres of ice crystal formation during vitrification, and the symbionts tolerated the trehalose sugar solution well. To further improve the vitrification, we flattened the sample to an ultra-thin alginate film on a modified cryotop blade by pre-wetting the blade in a solution of calcium chloride (used for the polymerisation of the alginate), which facilitated the breakdown of the surface tension of the alginate-encapsulated sample. We added one final step to ensure successful vitrification. After the first three equilibration steps, which remove most intracellular water and allow for the penetration of cryoprotectants, we dried the alginate-encapsulated sample using a Kimwipe, added a drop of the high-solute vitrification solution DPT (1M trehalose, 10% DMSO, 5% PG in PBS), which the sample was exposed to for 30 sec before being plunged into liquid nitrogen. Freshly extracted symbionts survived longer in culture-like conditions when they were well cleaned at the end of the extraction (i.e., with as few coral host cells remaining as possible), when they were exposed to antibiotics shortly after extraction, and when they were stored as droplets encapsulated in alginate. The same held for the recovery of symbionts after cryopreservation and laser warming. Therefore, during the extraction, we cleaned the symbionts to a point that they would easily pass through a 20-micrometer mesh filter, then exposed all symbionts to an antibiotic cocktail designed by Polne-Fuller (1) for a period of 12 h. Afterwards, they were rinsed three times through sequential spins in the centrifuge and resuspensions in filtered seawater and then passed through a 20-micrometer mesh filter one last time. During the post-thaw recovery, symbionts were left in their alginate films. This holding regime yielded better recovery and survival, as determined by pulse amplitude modulated fluorometry measurements and microscopic observations.

### Supplementary Information: Detailed laser-warming

Vitrified samples were warmed using a bench-top iWeld 980 Series infrared laser (LaserStar Technologies Corporation, Orlando, FL, USA). The laser was equipped with a stereomicroscope to allow visualisation of samples within the laser chamber. A custom-made cryo-jig (Design Solutions, Inc., Chanhassen, MN, USA) was used to lower the sample into liquid nitrogen for vitrification, raise the sample into the laser beam focal region, and trigger the laser for warming. The cryo-jig was positioned above a polystyrene bath containing liquid nitrogen, installed in the laser chamber. A control box with custom software (Design Solutions, Inc., Chanhassen, MN, USA) was used to control the position of the cryo-jig arm holding the cryostick and to fire the laser. For warming of vitrified symbionts, the control box was used in ‘AUTO’ mode so that a single laser pulse was fired automatically when the sample was raised into the laser beam focal region. Gold nanorods, as characterised by Khosla *et al*. (3), were used as a laser absorber for warming of vitrified samples. The nanorods (nanoComposix, San Diego, CA, USA) were first diluted with the algal symbionts in solution, before their encapsulation in alginate, at a ratio of 1:50, and also with the final vitrification solution, at a ratio of 1:100 for a final concentration of 70µg/mL with an optical density of 49 at the peak absorbance wavelength of 1064 nm throughout the sample and vitrification solution. This cryoprotectant solution containing gold nanorods was then used for vitrification and laser warming of *L. scutaria* symbionts.

For the laser warming, the final settings (330 V, 2 ms pulse width, and 2 mm laser spot diameter) were determined empirically by vitrifying our samples on the cryotops and observing the outcome when the droplet was warmed by the laser. The laser-warmed samples on cryotops were then immediately submerged into a solution containing 0.5M trehalose in FSW for rehydration and after 20 minutes, transferred to F2 medium (Sigma-Aldrich, G0154) in FSW and kept in the dark for 24 hrs to recover before viability assessments or use for coral larvae infections.

### Supplementary Information: Pulse amplitude modulated fluorometry

As part of their symbiosis with corals, algal symbionts provide essential photosynthetic products to their host. We used a Junior Pulse Amplitude Modulated fluorometer (Junior-PAM, Walz, Germany) to obtain an indication of the viability of photosynthesis in the Photosystem II. In dark-adapted conditions, the Pulse Amplitude Modulated fluorometer measures initial fluorescence (recorded as ‘$F_0$’), then administers a saturating pulse of light and records the maximal fluorescence (recorded as ‘$F_m$’). A ratio of these, $(F_m-F_0)/F_m$, gives the effective quantum yield, recorded as the Y-value (Y), which varies between 0 and 1. Settings of the Pulse Amplitude Modulated fluorometer, specifically the gain and intensity settings, were kept constant throughout each and across all experiments in these studies to ensure that we were observing physiological changes. The percent change from a sample’s initial quantum yield post-treatment was reported. For the purpose of this paper, any quantum yield or Y-value at or below 0.100 was defined as non-viable and as having a non-functional photosystem (4). By inference, quantum yields with the least change from their initial measures, indicate a robust and healthy algal symbiont sample, whereas those close to 100% change were mostly a dead population of cells. Measurements with the Pulse Amplitude Modulated fluorometer were conducted in dark conditions in the laboratory with photosynthetically active radiation values below 1 µmol photons $m^{-2}$ $s^{-1}$ (Apogee Instruments, Model MQ-200) for at least 30min before measurements. All solutions, including those containing the extracted symbionts, were prepared and maintained in 0.2 µm- filtered seawater, unless otherwise stated.

### Supplementary Information: Optimised protocol, step-by-step

The Symbiodiniaceae should be freshly isolated following Bouwmeester et al. (5), and prepared and encapsulated in 2% alginate following Bouwmeester et al. (6). Once the encapsulation is complete (after 30 minute polymerisation in the calcium chloride solution), the encapsulated Symbiodiniaceae will be exposed to different cryoprotectant solutions, in batches of 15 flattened droplets (3 cryoholders with 5 droplets each). See Bouwmeester et al. (6) regarding making cryoholders and cryoblades. All solution containing trehalose were made with trehalose from Pfanstiehl Inc. Waukehan, IL, USA, CAS # 6138-23-4.

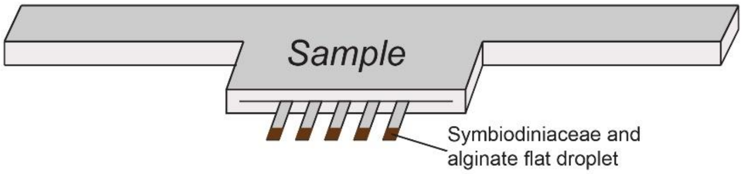


Figure 1: example of a cryoholder containing 5 acetate cryoblades. The encapsulated Symbiodiniaceae are then placed at the tip of the cryoblades.

1. Immerse the encapsulated samples in EQ1 (10% DMSO in FSW solution) for 60 minutes.

2. Immerse the encapsulated samples in EQ2 (0.5M trehalose in 10% DMSO in FSW solution) for 30 minutes.

For the next steps, there are two options, depending on whether the goal is to bank samples for long-term storage, or to rewarm them immediately.

**Option 1: Cryopreserving and banking samples for long-term storage**

For the following steps, proceed in batches of one cryoholder (and 5 samples) at a time.

3. Immerse the encapsulated samples in EQ3 (0.5M trehalose in 10% DMSO and 5% methanol in FSW solution) for 3 minutes.

4. Lift the cryoholder and its 5 blades out of EQ3, and dry the blades and the droplets as much as possible using a Kimwipe. This step is important as without it, the samples will not vitrify well.

5. Add a 2-µl droplet of DPT vitrification solution (1M trehalose, 10% DMSO, 5% PG, in PBS) containing gold nanorods, on the first sample, start a timer, and add 2-µl droplets of DPT on the remaining four samples.

6. At exactly 30 seconds on the timer, lower the cryoholder and blades into a liquid nitrogen bath, so that the bottom half of the cryoblades are immersed in the liquid nitrogen. The droplets should appear transparent, indicating that vitrification was successful. As much as possible, avoid submerging the cryoholder. If submerged at any time, it may be hard to pull the cryoblades out of the cryoholder.

7. Using a pre-cooled forceps, transfer each blade into a chilled 1.8-mL cryovial. The temperature inside the cryovial needs to be below -80°C for this step to be successful. We recommend using an aluminium block that can be placed directly into a liquid nitrogen bath, which will maintain the cryovials at the ideal temperature.

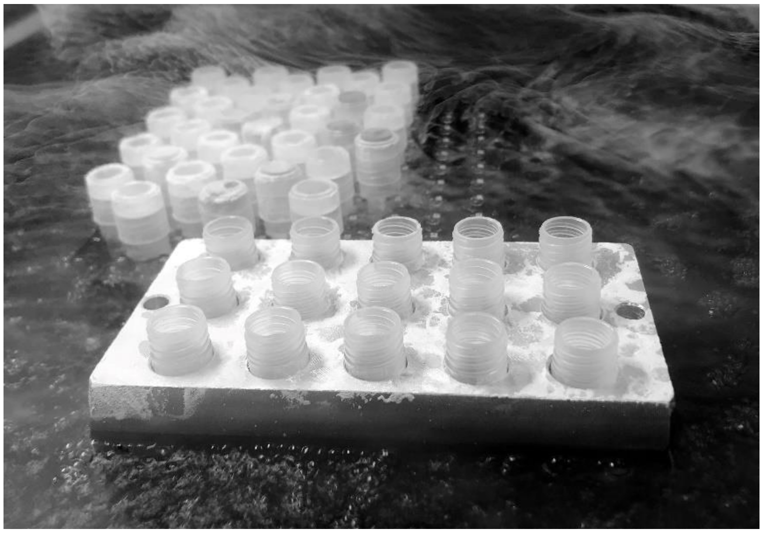

Figure 2: Aluminium block placed directly in a liquid nitrogen bath, holding the cryovials that will contain the vitrified Symbiodiniaceae

8. Once the cryovials are capped, transfer the cryovials into a cryobox. Remember to hold the box in a liquid nitrogen bath during this step.

9. Transfer the cryobox to a cryobank that will maintain the temperature of the samples at -196°C.

**Option 2: Cryopreserving samples for immediate re-warming and assessment**

Before starting, make sure that the infrared laser is on, the laser jig connected, and that the laser liquid nitrogen bath is in place. We use a bench-top iWeld 980 Series infrared laser (LaserStar Technologies Corporation, Orlando, FL, USA) equipped with a stereomicroscope to allow visualisation of samples within the laser chamber and a custom-made cryo-jig (Design Solutions, Inc., Chanhassen, MN, USA) that was used to (1) lower the sample into liquid nitrogen for vitrification, (2) raise the sample into the laser beam focal region, and (3) trigger the laser for warming. The cryo-jig was positioned above a polystyrene bath containing liquid nitrogen, installed in the laser chamber. A control box with custom software (Design Solutions, Inc., Chanhassen, MN, USA) was used to control the position of the cryo-jig arm holding the cryotop and to fire the laser. Set the laser settings on 330 V, 2mS, 0 Hz, 2 mm diameter, flat profile.

For the following steps, proceed one sample at a time.

3. Using a forceps, transfer one blade from its cryoholder on to a cryostick

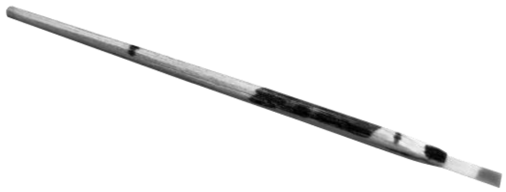

Figure 3: Cryostick, made from a bamboo barbecue skewer, tapered at one end to fit in the laser-warming jig (3) and adjusted to fit a cryoblade at the other end. Markings are added on the cryostick to quickly identify which side the sample is facing on the cryoblade, to know how far to insert the cryostick in the laser jig and to know how far to insert the cryoblades in the cryostick.

4. Immerse the encapsulated samples in EQ3 (0.5M trehalose in 10% DMSO and 5% methanol in FSW solution) for 3 minutes.

5. Lift the cryostick and blade out of EQ3, and dry the blade and droplet as much as possible using a Kimwipe. This step is important as without it, the sample will not vitrify well.

6. Transfer the cryostick into the laser chamber and insert the end of the cryostick into the laser jig.

7. Add a 2-µl droplet of DPT vitrification solution (1M trehalose, 10% DMSO, 5% PG, in PBS) containing gold nanorods, on the sample and start a timer.

8. At exactly 30 seconds on the timer, lower the cryostick and blade into a liquid nitrogen bath using the laser jig control box, so that the bottom half of the cryoblade is immersed in the liquid nitrogen.

9. Wait 10-20 seconds until the sample has stopped boiling in the liquid nitrogen, indicating that the sample's temperature has reached liquid nitrogen temperature.

10. Raise the laser jig arm using the laser jig control box and fire the infrared laser (laser settings: 330 V, 2mS, 0 Hz, 2 mm diameter, flat profile)

11. Remove the cryostick from the laser chamber and immediately plunge the sample in a rehydration solution of 0.5 M trehalose in FSW

12. Transfer the cryoblade to a cryoholder sitting above a basin containing a 0.5 M trehalose in FSW rehydration solution and start a timer

13. After 20 minutes, transfer the cryoblade to a new cryoholder sitting above a new basin with F/2 marine water enrichment solution (Sigma-Aldrich, G0154).

14. Once all cryoblades have been retrieved from laser-warming, move the F/2 marine water basin containing all cryoblades and samples to an incubator set at ambient seawater temperature (25-26 °C for Hawaiian Symbiodiniaceae) and cover the container with aluminium foil to prevent any photosynthetic activity of the algal symbionts, to facilitate recovery

15. The next day, the photosynthetic activity of the Symbiodiniaceae should be assessed using a Pulse Amplitude Modulated fluorometer (e.g., Junior-PAM, Walz, Germany) and should be compared to PAM-assessments from control samples that have not undergone cryopreservation.

### Supplementary Information: References

### Supplementary Figure

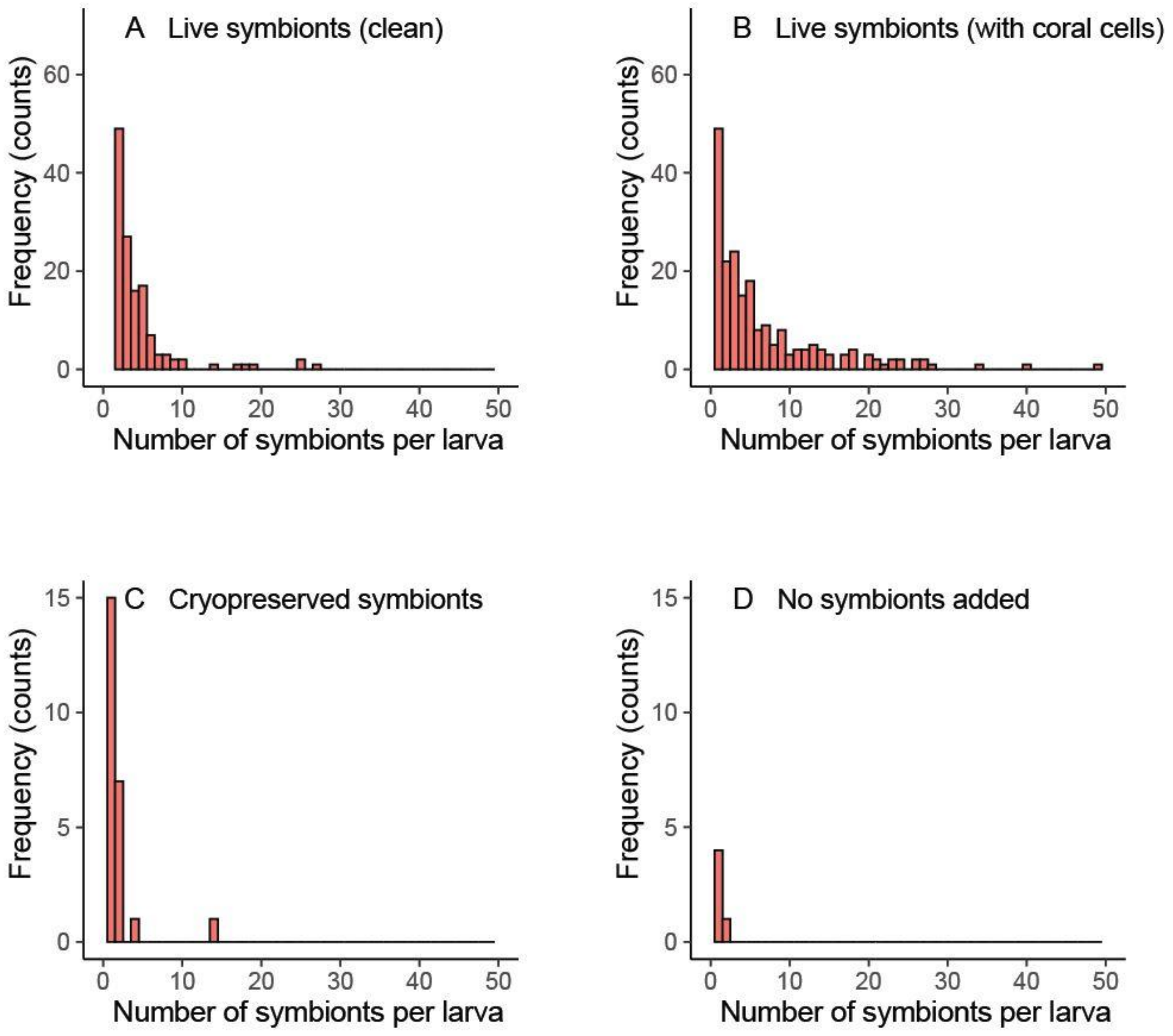


**Figure S1: Frequency distribution of the number of symbionts taken up per larva** in the clean fresh symbiont (A), fresh symbiont with host coral cells (B), cryopreserved symbiont (C), and no symbiont added (D) treatments.

### Supplementary Tables (S1 – S3)

**Table S1: Summary of ANOVA results** and related posthoc tests on the photosynthetic yield of algal symbionts after treatment (Cryopreserved, Toxicity, Live, or Dead)

A. Algal symbionts extracted from *Lobactis scutaria*

| ***One-way ANOVA*** | | | | |
|---|---|---|---|---|
| **Sum of Squares** | **DF** | **F-value** | **p-value** | |
| 0.469 | 3 | 21.39 | $6 \times 10^{-7}$ | *** |
| ***Least square means posthoc test with Tukey adjustment for multiple comparisons*** | | | | |
| **Treatment** | **Least square mean** | **SE** | **DF** | **group** |
| Cryo | 0.313 | 0.024 | 24 | b |
| Tox | 0.363 | 0.038 | 24 | b |
| Live | 0.431 | 0.038 | 24 | b |
| Dead | 0.031 | 0.038 | 24 | a |

B. Algal symbionts extracted from *Porites compressa*

| ***One-way ANOVA*** | | | | |
|---|---|---|---|---|
| **Sum of Squares** | **DF** | **F-value** | **p-value** | |
| 0.304 | 3 | 31.56 | $3 \times 10^{-8}$ | *** |

| ***Least square means posthoc test with Tukey adjustment for multiple comparisons*** | | | | |
|---|---|---|---|---|
| **Treatment** | **Least square mean** | **SE** | **DF** | **group** |
| Cryo | 0.220 | 0.014 | 23 | b |
| Tox | 0.178 | 0.033 | 23 | b |
| Live | 0.439 | 0.028 | 23 | c |
| Dead | 0.030 | 0.033 | 23 | a |

C. Algal symbionts extracted from *Leptastrea purpurea*

| ***One-way ANOVA*** | | | | |
|---|---|---|---|---|
| **Sum of Squares** | **DF** | **F-value** | **p-value** | |
| 0.705 | 3 | 535.63 | $2 \times 10^{-16}$ | *** |

| ***Least square means posthoc test with Tukey adjustment for multiple comparisons*** | | | | |
|---|---|---|---|---|
| **Treatment** | **Least square mean** | **SE** | **DF** | **group** |
| Cryo | 0.134 | 0.009 | 17 | b |
| Tox | 0.419 | 0.009 | 17 | c |
| Live | 0.435 | 0.009 | 17 | c |
| Dead | 0.017 | 0.009 | 17 | a |

**Table S2: Photosynthetic yield summary of algal symbionts used for symbiont uptake by coral larvae experiments**, determined with a pulse amplitude modulated fluorometer (Junior-PAM, Walz, Germany)

| Treatment | Time between preparation and use for experiments | Photosynthetic Yield* (mean ± SEM) |
|---|---|---|
| Live with Coral Cells, in solution | 1 h | 0.600 ± 0.014 |
| Live Clean, in alginate | 2 days | 0.439 ± 0.006 |
| Cryopreserved, in alginate | 1 day | 0.313 ± 0.031 |
| Dead, in alginate, after all cryopreservation steps but the laser warming | 1 day | 0.018 ± 0.003 |
| Dead, in solution, after multiple freeze-thaw cycles | 1 h | 0.000 ± 0.000 |

* Measured at the time of use for experiments

**Table S3: Summary of Logistic Regression results** and related posthoc tests on the algal symbiont uptake success by *L. scutaria* larvae in each algal symbiont treatment (Freshly isolated algae, Cryopreserved algae, No algae added, or Dead algae)

| *Logistic Regression on Freshly isolated algae treatments: unfiltered vs clean* | | | |
|---|---|---|---|
| **Wald's $\chi^2$** | **DF** | **p-value** | |
| 139.58 | 1 | $<2 \times 10^{-16}$ | *** |
| ***Logistic Regression on Cryopreserved symbiont concentrations: 1.5 × 10$^5$, 7.5 × 10$^4$, 3 × 10$^4$ and 1.5 × 10$^4$ cells/mL*** | | | |
| **Wald's $\chi^2$** | **DF** | **p-value** | |
| 0.82 | 3 | 0.846 | |
| ***Logistic Regression on Dead symbiont treatments: vitrified and ambiently warmed (without laser) versus exposed to multiple cycles of freeze-thaw*** | | | |
| **Wald's $\chi^2$** | **DF** | **p-value** | |
| 1.017 | 1 | 0.313 | |
| ***Logistic Regression on Cryopreserved versus Dead and No Symbionts added treatments*** | | | |
| **Wald's $\chi^2$** | **DF** | **p-value** | |
| 7.85 | 2 | 0.020 | * |
| ***Logistic Regression posthoc contrasts with Sidak adjustment for multiple comparisons*** | | | |
| **Treatment Effect** | **p-value** | | |
| Cryopreserved algae | 0.021 | * | |
| No algae added | 0.726 | | |
| Dead algae | 0.720 | | |
